# Gene pseudogenization in fertility-associated genes in cheetah (*Acinonyx jubatus*), a species with long-term low effective population size

**DOI:** 10.1101/2024.04.26.591110

**Authors:** Jessica A. Peers, Will J. Nash, Wilfried Haerty

**Affiliations:** Earlham Institute, Norwich Research Park, UK

**Author notes:** Corresponding author, Norwich Research Park, Colney Ln, Norwich NR4 7UZ. These authors contributed equally (Will J. Nash, Wilfried Haerty).

**Keywords:** pseudogenization, male fertility, cheetah, inbreeding, effective population size, premature termination codons

## Abstract

The ongoing global biodiversity crisis is placing an increasing number of mammalian populations at risk of decline. Species that have survived severe historic bottlenecks, such as the cheetah (*Acinonyx jubatus*) exhibit symptoms of inbreeding depression including reproductive and developmental defects. Although it has long been suggested that such defects stem from an accumulation of weakly deleterious mutations, the implications of such mutations leading to pseudogenization has not been assessed.

Here, we use comparative analysis of eight felid genomes to better understand the impacts of deleterious mutations in the cheetah. We find novel pseudogenization events specific to the cheetah. Through careful curation, we identify 89 genes with previously unreported premature termination codons that likely affect gene function, 65 of which are caused by point mutations. With the addition of population data, we find 22 PTCs fixed in wild populations, four of which (DEFB116, ARL13A, CFAP119 and NC5TD4) are also found in a more recent reference genome. Mutations within three of these genes are linked with sterility, including azoospermia, which is common in cheetahs. Our results highlight the power of comparative genomic approaches for the discovery of novel causative variants in declining species.

## Introduction

The ongoing biodiversity crisis is placing an increasing number of mammalian species at risk of population decline (Ceballos and Ehrlich, 2023). Lower effective population size (*N_e_*) leads weakly deleterious alleles to behave ‘nearly neutrally’, enabling them to segregate at high frequencies (Crow and Kimura, 1970; Kimura, 1983; Ohta, 1992; Björnerfeldt *et al*., 2006). The accumulation of such deleterious mutations negatively impacts fitness and can lead to genetic inviability placing declining populations in an even more precarious position (Lynch *et al*., 1993; Lande, 1994).

In addition to missense mutations that lead amino acid replacement, deleterious mutations can occur in the form of nonsense or frameshift mutations in protein-coding genes, potentially resulting in the loss of coding potential due to the creation of premature termination codons (PTCs). PTCs can lead to exon skipping, decreased mRNA stability and protein truncation, resulting in the transcripts being targeted by the nonsense mediated decay pathway (Khajavi *et al*., 2006). The resultant sequences are known as pseudogenes; vestigial coding sequences that remain in the genome but, due to deleterious mutations, are not successfully translated (Li *et al*., 1981).

Pseudogenization is predominantly observed in genes with little contribution to fitness, where deleterious mutations can accumulate in the absence of selection <u>(Ochman and Davalos, 2006)</u>. However, in the case of small, inbred populations, as weakly deleterious mutations accumulate at a higher rate than they would in larger populations, genes with greater contributions to an organism’s fitness can become pseudogenized <u>(Ochman and Davalos, 2006)</u>.

Although many studies have considered the impacts of functionally-damaging missense variants in species with low *N_e_* (Daetwyler *et al*., 2014; Dobrynin *et al*., 2015; Marsden *et al*., 2016; Samaha *et al*., 2021), elevated rate of allele loss <u>(Masel, 2011)</u>, and increased likelihood of homozygous expression of a recessive deleterious allele (Mukai *et al*., 1972; Charlesworth and Charlesworth, 1987; Charlesworth and Willis, 2009; Bosse *et al*., 2019), pseudogenization as a result of population decline has often been overlooked in the literature.

To address this gap, we focus on the cheetah, *Acinonyx jubatus*, to investigate the effects of long-term low *N_e_* on patterns of pseudogenisation. The cheetah has been suggested to have exhibited a much lower *N_e_* over the last 3 million years compared to other felid species (Kim *et al*., 2016). When the cheetah migrated put of the Americas and into Eurasia/Africa over 100,000 years ago, a genetic bottleneck occurred (Dobrynin *et al*., 2015). Additionally, cheetah populations experienced a severe bottleneck approximately 10-12,000 years ago at the end of the Pleistocene epoch, suggested to be the origin of the current lack of genetic diversity of the species (O’Brien *et al*., 1987; Menotti-Raymond and O’Brien, 1993; Driscoll *et al*., 2002; Dobrynin *et al*., 2015; Terrell *et al*., 2016). Finally, poaching, habitat loss, prey loss, and human-wildlife conflict all impact current cheetah populations, leading them to be classified ‘Vulnerable’ by the IUCN <u>(Durant, S.M.,</u> <u>Groom, R., Ipavec, A., Mitchell, N. & Khalatbari, L, 2021)</u>. Contemporary population size estimates remain low, with an estimated census size of ∼7,100 (Marker *et al*., 2008; Durant *et al*., 2017).

Population genetic studies have shown the cheetah to have experienced prolonged inbreeding to the point that there is no recorded non-inbred wild population <u>(O’Brien and Johnson, 2005)</u>. This led to high homozygosity, accumulation of weakly deleterious mutations, and severe inbreeding depression (Dobrynin *et al*., 2015). Cheetahs exhibit increased susceptibility to infectious diseases (O’Brien *et al*., 1985; Terio *et al*., 2018), developmental instability (Wayne *et al*., 1986), a high frequency of spermatozoal abnormalities (Wildt *et al*., 1983) and high juvenile mortality in the wild and in captivity (O’Brien *et al*., 1985; Bell, 2005). High levels of homozygosity result in a lack of diversity at the MHC loci enabling viable skin grafts to be made between unrelated cheetahs (O’Brien *et al*., 1985). Putatively deleterious non-synonymous mutations have been identified in genes with functions related to observed abnormalities in cheetah populations (Dobrynin *et al*., 2015; Samaha *et al*., 2021). However, these studies did not explore loss of function due to premature termination codons (PTC), despite the potential for pseudogenization to also contribute to the observed morphological and physiological defects.

Here, we investigate pseudogenization in the cheetah by taking advantage of the increased availability of genomic resources to compare the reference genomes of eight felid species: cheetah (*Acinonyx jubatus*), domestic cat (*Felis catus*), black-footed cat (*F. nigripes*), lion (*Panthera leo*), jaguar (*P. onca*), leopard (*P. pardus*), tiger (*P. tigris*) and puma (*Puma concolor*) (Armstrong *et al*., 2020; Zoonomia Consortium, 2020; Christmas *et al*., 2023). After careful curation, we identify 89 genes with novel cheetah-specific PTCs. In four of these genes (ARL13A, DEFB116, CFAP119, NC5TD4), potentially involved in reproduction and susceptibility to infectious diseases, issues prevalent in wild cheetah populations, we find support for these novel cheetah-specific PTCs in multiple unrelated cheetahs. Pseudogenisation of these genes could be of interest for conservation and this finding contributes to our understanding of the effect of long-term low *Ne* on the genome.

## Materials and Methods

### Gene family identification

Publicly available data for twelve mammal species were used in this study (Table S1): eight felid species and four mammal outgroups. Coding sequences (CDS) were extracted from each of ten species using annotations generated in Christmas et al. (2023) (Table S1). CDS files for Human (GCA_000001405.15) and Mouse (GCA_000001635.2) were downloaded from Ensembl release 99 (Cunningham *et al*., 2022) (Table S1). The longest transcript (nucleotide length) per gene was selected for analysis (https://github.com/TGAC/GSTF_snakemake). A species tree was inferred using topologies from existing literature (Piras *et al*., 2018; Armstrong *et al*., 2020; Zoonomia Consortium, 2020) (see Figure 1). Gene families were then clustered into gene trees using GeneSeqToFamily (GSTF) (Thanki et al., 2018; https://github.com/TGAC/GSTF_snakemake).

**Figure 1.**
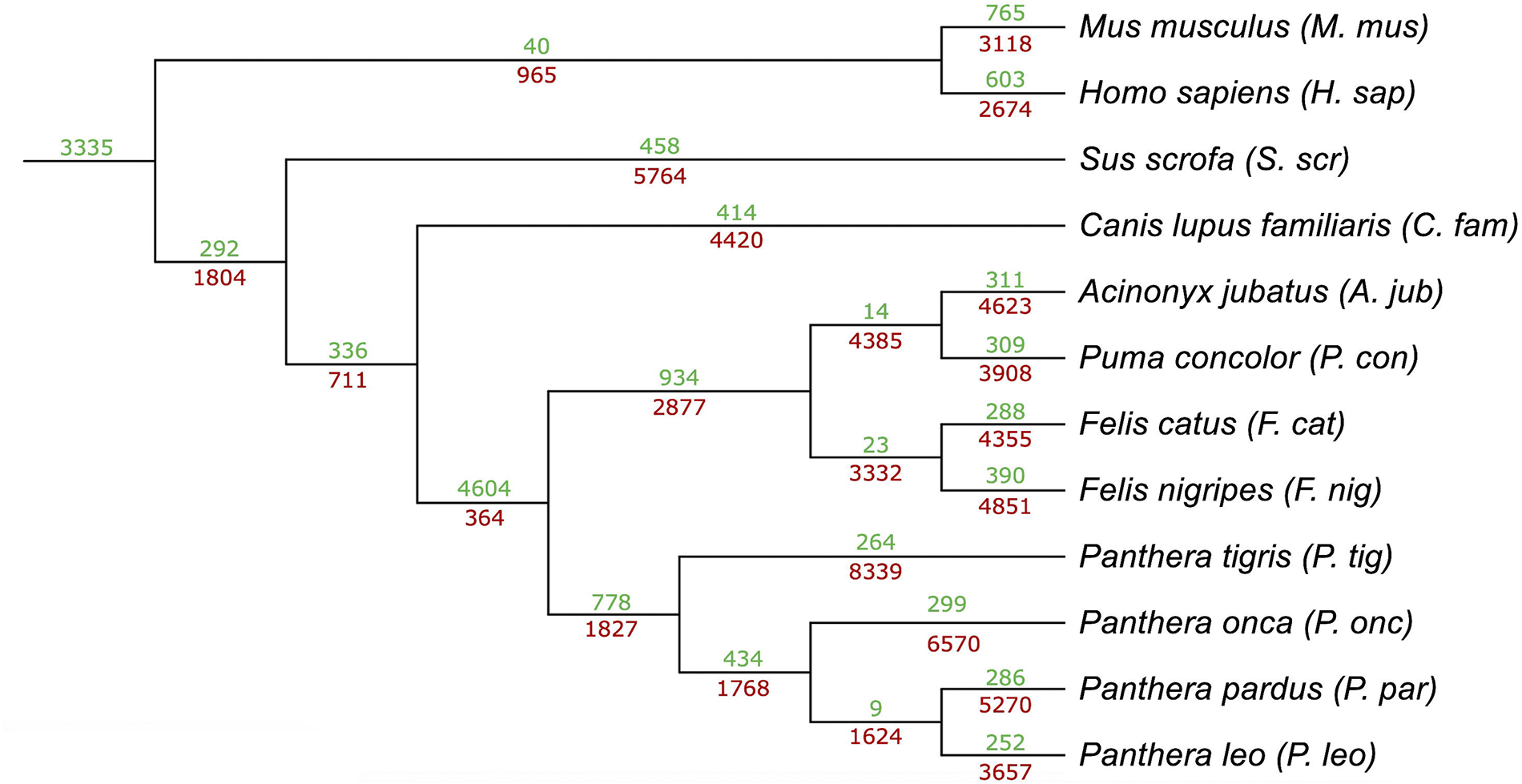
Gene gains and losses in the Felidae. *NOTUNG* (Chen *et al*., 2000), a tree reconciliation tool, was run on the gene trees generated by *GeneSeqToFamily* (Thanki *et al*., 2018). Gene gains and losses identified by *NOTUNG* for each branch of the tree are shown in green and red, respectively. The topology of the species tree was extracted from existing literature (Piras *et al*., 2018; Armstrong *et al*., 2020; Zoonomia Consortium, 2020).

### Gene tree reconciliation

To identify genes lost in the cheetah, tree reconciliation was conducted on gene trees generated by *GSTF* using *NOTUNG* v.3.0.26 (Chen *et al*., 2000), resulting in quantification of gene gains and losses at each node in the species tree. *NOTUNG* does not report a specific gene loss and rather reports a loss in a gene tree. Not every gene tree contains exactly one gene per species, so it was necessary to interrogate each tree with a reported loss to identify the missing gene.

### Curation of results

Each gene with a reported loss in the cheetah was further investigated to determine presence or absence in the aciJub1 annotation (GCA_001443585.1).

### Genes present in aciJub1 annotation

Genes in the cheetah annotation which did not follow coding logic (length not a multiple of three, premature termination codon) were filtered out by GSTF (Thanki et al., 2018; https://github.com/TGAC/GSTF_snakemake). To determine whether such genes contain biologically feasible premature termination codons (as opposed to misannotation or sequencing errors), orthologous puma (*Puma concolor, PumCon1.0, GCA_003327715.1*) sequences were used as a closely related reference.

Stringent filtering was then undertaken to identify the most biologically feasible mutations. Genes with no annotated copy in the puma were filtered out. For each remaining gene, the difference in nucleotide length between the cheetah and puma sequences was calculated and genes with a difference in length of at least 10% were filtered out. Nucleotide sequences were translated to protein (*EMBOSS* v6.6.0 *Transeq,* (Rice *et al*., 2000) and sequence identity between cheetah and puma sequences was calculated using the Smith-Waterman algorithm (*EMBOSS* v6.6.0 *Water,* (Rice *et al*., 2000). Genes with a sequence identity of below 90 % were removed due to potential misannotation and genes with a sequence identity of 100 % were removed as a PTC is expected to be one of multiple mutations accumulating in a sequence; therefore, PTCs in an otherwise identical sequence are more likely to be sequencing errors. Following translation, amino acid sequences corresponding to each transcript were used to identify PTCs. Where no PTC was identified, the transcript was filtered out.

Of the remaining genes, highly-duplicated or fast-evolving gene families (Zinc-finger proteins (ZNF), olfactory receptors (OR), family with sequence similarity (FAM), transmembrane proteins (TMEM), cyclin-dependent kinases (CDK) and cytochrome P genes (CYP)) were filtered out, as pseudogenization is expected in these gene families (Nei *et al*., 2008; Albà, 2017). Transcripts retired from the current ENSEMBL annotation were also filtered out at this stage.

For the remaining genes, an amino acid sequence alignment was generated by adding the corresponding aciJub1 (GCA_001443585.1) sequence to the *GSTF* gene family alignment and realigning (*MAFFT* v.2.071, <u>(Katoh and Standley, 2013)</u>, default settings. Through careful manual curation of alignments, the remaining genes were classified based on whether the PTC is unique to the cheetah (versus shared with the puma lineage) and whether the PTC is due to mis-annotation or a biologically feasible frameshift or point mutation. The sequence identity of regions upstream and downstream (excluding a buffer of 30 nucleotides either side to minimise the impact of frameshifts) of putative frameshifts and point mutations was calculated using the Smith-Waterman algorithm (*EMBOSS* v6.6.0 *Water,* (Rice *et al*., 2000) and genes with a sequence identity of over 97 % were classified as less likely to be biologically feasible.

During the course of this study, a novel long-read cheetah assembly (VMU_Ajub_asm_v1.0, GCA_027475565.2) was published (Winter *et al*., 2023). CDS sequences from this genome were added to PTC candidate alignments to identify if the biologically feasible PTCs are likely to be specific to the individual sequenced for the aciJub1 reference genome or if they are potentially present across the species.

### Genes not present in aciJub1 annotation

Putative gene losses in the cheetah may also represent annotation or assembly errors. Genes with no annotated copy in the puma were filtered out as these gene losses may have occurred earlier in the cheetah/puma lineage. The cheetah genome was searched for evidence of unannotated genes, using puma CDS sequences (*blastn, BLAST* v2.10 (Camacho *et al*., 2009). *BLAST* hits were filtered to retain only those with e-value < 10^-6^, or e-value < 10^-3^ if the length of the hit was < 50 nt. *BLAST* hits that overlapped with annotated regions of the cheetah genome were filtered out. Successful hits were reciprocally *BLASTed* to the puma genome to ensure that *BLAST* hits represented all exons of the gene.

Exonic sequences of genes that had not been identified in the aciJub1 assembly were then searched for individually using *BLAT* v.35 <u>(Kent,</u> <u>2002)</u>to identify genes where at least one exon was missing, a PTC was present, or part of the gene sequence overlapped with another gene, resulting in misannotation in the cheetah. Highly conserved synteny between the cheetah and puma (Armstrong *et al*., 2020) was exploited to determine whether the *BLAST/BLAT* hits were located in the expected region of the genome. For genes with no significant *BLAST/BLAT* hit, this synteny was utilised to reject genes likely missing due to high fragmentation of the assembly.

### Population analyses

Low coverage whole genome resequencing data for 6 cheetahs (Dobrynin et al., 2015) was downloaded from Genbank (SRR2737543-SRR2737545). Reads were trimmed using *Trimmomatic* v.039 (Bolger, Lohse and Usadel, 2014) and mapped to the aciJub1 (GCA_001443585.1) reference using *BWA-MEM* v0.7.17 (Li, 2013). Mapping was followed by *SAMtools* v1.15 *fixmate* and *sort* (Danecek et al., 2021). PCR duplicates were then removed using *Picard* v2.26.2 *RemoveDuplicates* (Broad Institute, 2019). Finally, mappings were filtered for complete read pairs and mappings of MAPQ >25 using samtools view (see Table S2 for settings).

Joint genotyping was conducted using *BCFtools* v1.10.2 *mpileup* (Danecek *et al*., 2021). *BCFtools call* was then used to call multi-allelic variants. Variants were then filtered using *BCFtools filter* and single nucleotide polymorphisms (SNPs) were extracted using *BCFtools view*. This generated a set of SNPs which were not within 3bp of other variants, had a variant quality score >= 30, that were at a locus with sequencing depth greater than 12 and less than 106 (+/-3 times average sequencing depth), had a minor allele count of 3 or more, and were represented by data at that locus in more than 50% of individuals.

SNPs were then intersected with the genomic coordinates of predicted PTCs using *bedtools* v2.30.0 *intersect* <u>(Quinlan and Hall, 2010)</u>. *bedtools* v2.30.0 *coverage* <u>(Quinlan and Hall, 2010)</u> was used to identify candidates without enough coverage to call SNPs. SNPs that intersected with predicted PTC coordinates were manually assessed to determine the prevalence of each PTC in the population data. Predicted PTC sites that were not reported as SNPs in the population data were interrogated to determine if the site was filtered out due to low coverage or if the site was not reported as all individuals matched the reference (and therefore the PTC was identified in all individuals).

## Results

### Gene family identification

From 230,740 coding sequences (Table S3), 9,673 CDS were filtered out as they did not follow coding logic, resulting in 221,067 CDS for which the longest nucleotide transcript was extracted (Table S3).

Following an internal filtering step (Table S3), GSTF sorted 207,314 transcripts into 13,979 clusters. Of these, 2,742 clusters with less than 3 genes were filtered out. The remaining clusters were split then into 19,349 gene trees. 6,671 1:1 orthologs were identified across all species, and 9,010 1:1 orthologs were present specifically in Felidae. Following gene tree reconciliation, gene gains and losses were identified across the dataset (Figure 1).

### Computational validation of results

4,623 gene trees with a putative gene loss in the cheetah were extracted (Table S4). Within these gene trees, 2,477 genes were absent from the expected orthogroup. 2,093 genes were subsequently characterised as putative gene losses as they were not present in any gene tree. 1,938 of these were identified as annotated aciJub1 transcripts. Following *BLAST* and *BLAT* searches, none of the remaining 155 unannotated genes were found to be pseudogenisation candidates (Table S5).

### Putative gene losses present in aciJub1 annotation

Of the 1,938 annotated transcripts not found in gene trees, following comparison to the puma genome and stringent filtering, 370 putative gene losses remained (Tables S6 & S7). 51 of these genes were filtered out as they are part of highly duplicated or fast-evolving gene families (16 ZNF, 13 OR, 3 FAM, 2 TMEM, 2 CDK, 1 CYP) or did not have a known gene name annotated (11 genes), leaving 318 genes.

Of the 318 remaining pseudogenisation candidates, 19 genes had a PTC in both the cheetah and puma (Table S8). Following manual assessment of nucleotide and amino acid alignments, one of these candidates (NC5TD4) was retained as a putative cheetah-specific pseudogenisation event.

The remaining 299 genes did not have PTCs identified in the puma and are candidates for cheetah-specific pseudogenisation (Table S9). Through careful manual curation of the coding sequence alignments, a further 88 genes were identified with a novel PTC unique to the cheetah (Table S9).

Of the 89 total pseudogenization candidates, 62 were due to point mutations, 24 were due to frameshift mutations and 3 had occurrences of both (Table S10). Of these, 4 PTCs are shared between aciJub1 and VMU_Ajub_asm_v1.0 assemblies: DEFB116, ARL13A, NC5TD4 (Figure 2) and CFAP119.

**Figure 2.**
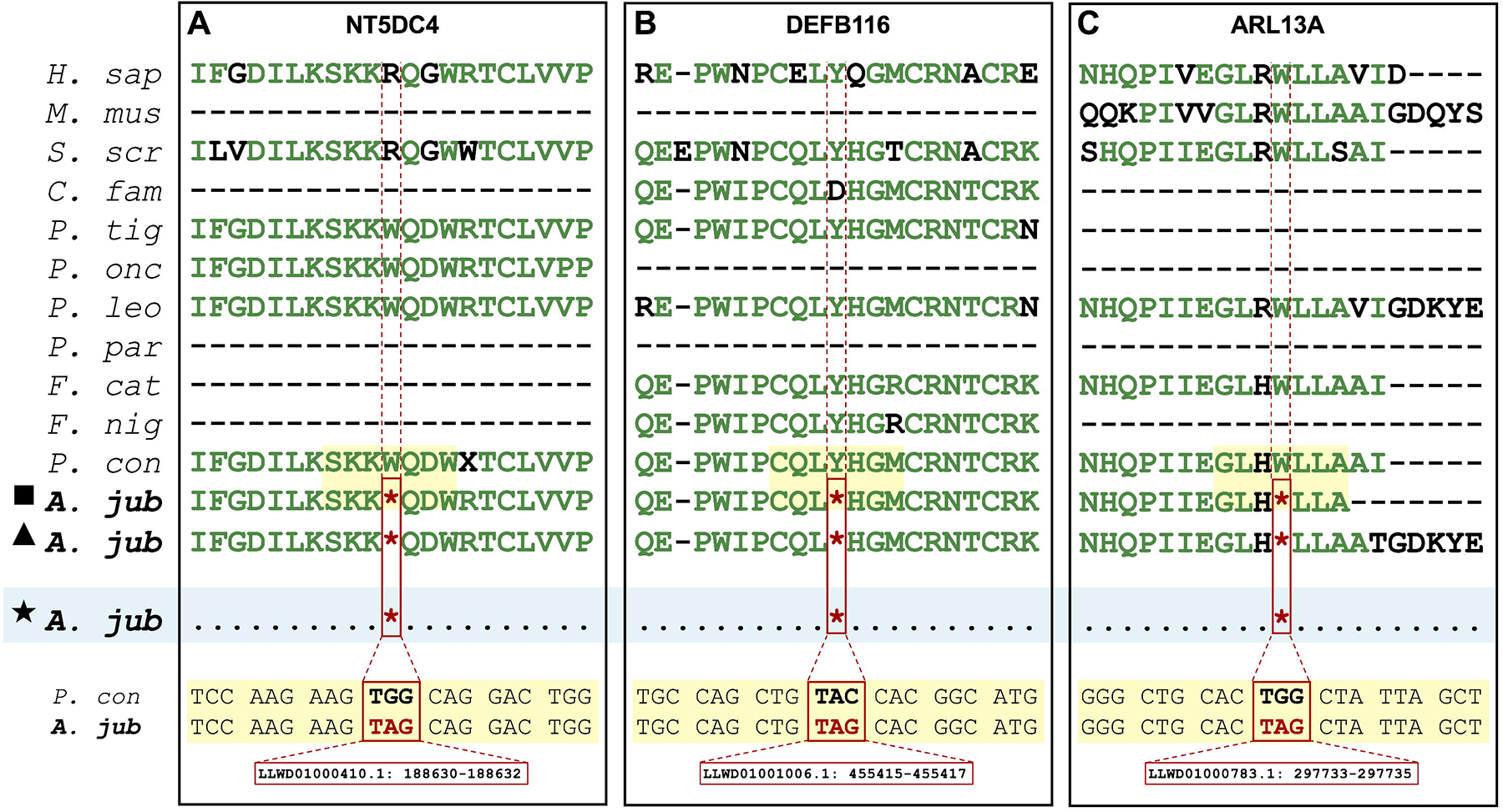
Putative pseudogenisation events in the aciJub1 cheetah genome. (PTCs) identified in ARL13A, NT5DC4 and DEFB116 in aciJub1 (▄), VMU_Ajub_asm_v1.0 (▴) and six African individuals (★) (Dobrynin *et al*., 2015). A protein alignment for each gene is shown for all species (expect those where the gene is unannotated, shown with dashes (-)). Species names correspond to those in Figure 1. Conserved amino acids are shown in green. The genomic position of each PTC is shown below the nucleotide sequence.

### Population genomic analysis of PTCs

Population data for six individuals (Dobrynin *et al*., 2015) was mapped to the aciJub1 genome and variants were called. Following conservative filtering (Table S2) 3,894,283 variant sites were identified, from which 1,572,165 high confidence SNPs were retained.

None of the 24 genes with PTCs caused by frameshift mutations had sufficient population resequencing coverage to determine the prevalence of the frameshift mutation in the population (Table S10). Of the 65 genes with PTCs caused by point mutations in aciJub1 (Table S10), 19 genes containing PTCs did not have sufficient population resequencing coverage to confidently determine the alleles in the population. A total of 22 PTCs were identified in all individuals suggesting those PTCs are fixed in those populations. These include the four previously identified pseudogenisation candidates (DEFB116, ARL13A, CFAP119 and NC5TD4). The PTC identified in CFAP119 occurs upstream of one of two protein domains implicated in a previously identified pseudogenisation event in Nordic Red dairy cattle (Figure 3, Iso-Touro et al., 2019).

**Figure 3.**
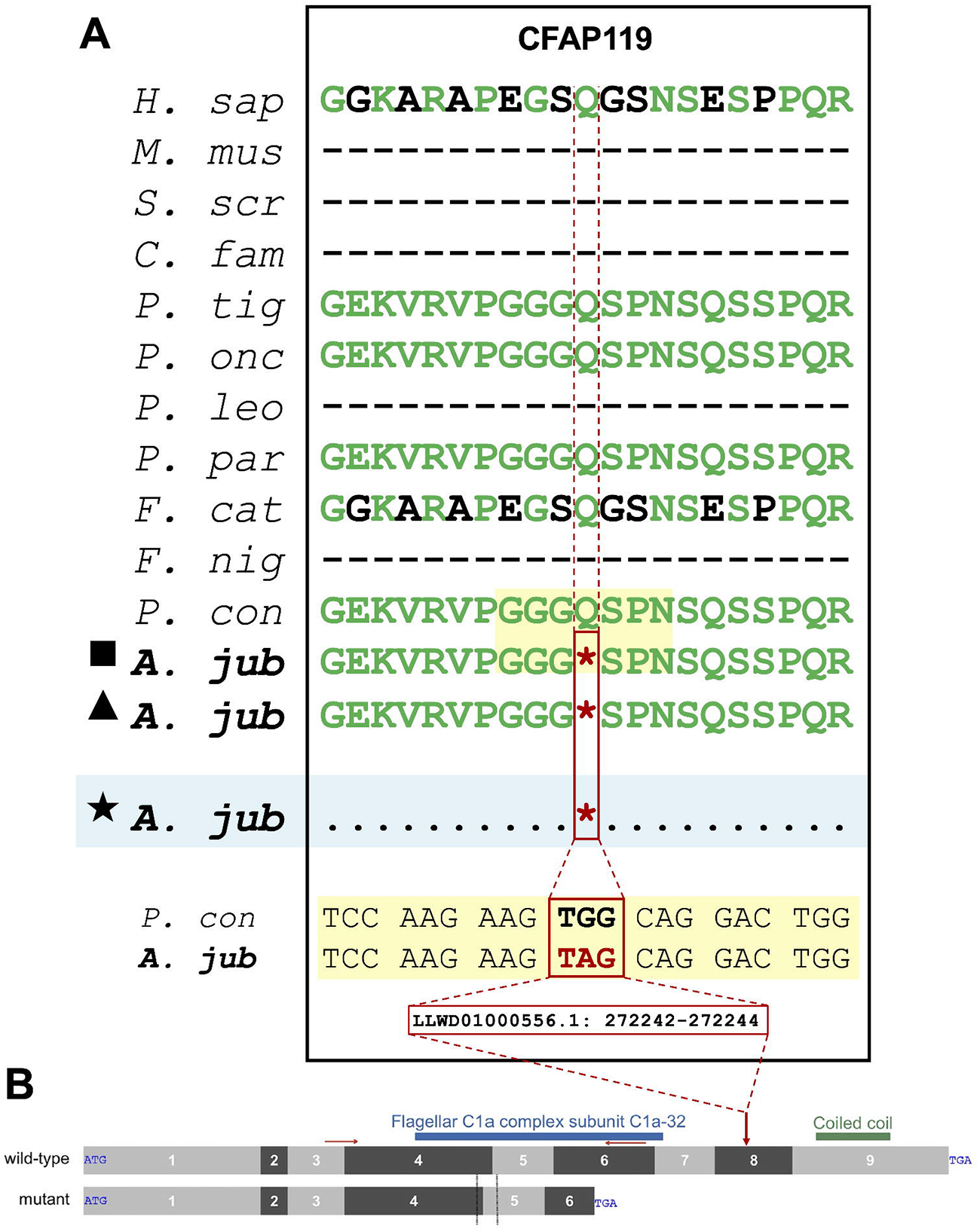
A novel cheetah-specific premature termination codon in CFAP119 has direct similarity to a fertility relevant pseudogenisation event in cattle. (A) Novel cheetah-specific premature termination codon (PTCs) identified in CFAP119 in aciJub1 (▄), VMU_Ajub_asm_v1.0 (▴) and six African individuals (★) (Dobrynin *et al*., 2015). See Figure 2 for formatting. (B) A splice donor variant in CFAP119 is associated in asthenospermia in Nordic red dairy cattle (extracted from Figure 3B&C in Iso-Touru et al. (2019)). Approximate position of the cheetah PTC on the protein is shown with a red arrow.

## Discussion

Historically bottlenecked populations, such as the cheetah, are characterised by low *Ne* and high inbreeding. This has been shown to have led to an accumulation of deleterious mutations (Dobrynin *et al*., 2015), with the potential to cause gene pseudogenization. In this study, we utilised the contemporary genomic data to compare felid genomes and identify cheetah-specific gene pseudogenization due to premature termination codons. Using coding sequences from eight felid species and four mammalian outgroups we annotated gene families and used tree reconciliation to identify cheetah-specific gene losses. Following validation of results, 89 genes with premature termination codons specific to the aciJub1 assembly were identified, including four genes (DEFB116, ARL13A, CFAP119, NC5TD4) with PTCs shared between all individuals investigated in this study, which may be related to reproduction and infectious disease susceptibility, known issues in the cheetah.

### Premature termination codons shared between multiple wild cheetahs

Of the 65 genes with biologically feasible premature termination codons in the original reference genome (aciJub1) caused by point mutations, 22 were fixed in population data generated by (Dobrynin *et al*., 2015) (Table S10).

Notable genes amongst these 22 include eight associated with male fertility (MOV10L1 (Fu *et al*., 2016), PHF7 (Cheng *et al*., 2023), ABHD10 (Smith and Eppig, 2009), CFAP119 (Iso-Touru *et al*., 2019), MARCHF6 (Smith and Eppig, 2009), MAGEB4 (Okutman *et al*., 2017), DEFB116 (Caballero-Campo *et al*., 2014; Zhang *et al*., 2018), and ARL13A (Schürmann *et al*., 2002)); immunity (DEFB116 (Schröder and Harder, 1999; Schneider *et al*., 2005; Dhople *et al*., 2006), ARL13A <u>(Song and</u> <u>Perkins, 2018)</u>, and IGBP1C (Smith and Eppig, 2009)); cancer (SSX5 (Smith and McNeel, 2010), TNFAIP1 (Tian *et al*., 2015), PRSS3 (Hockla *et al*., 2012), SLC38A7 (Haratake *et al*., 2021), and HAS3 (Wang *et al*., 2022)); and developmental defects (NCDN (Fatima *et al*., 2021)).

It has long been recognised that cheetah populations suffer defects in male fertility (Wildt *et al*., 1983), immunity (O’Brien *et al*., 1985; Terio *et al*., 2018) and development (Wayne *et al*., 1986). Additionally, cancer rates in wild and captive-managed felids have been shown to be high (Moresco *et al*., 2020), potentially due to the high fat and low fibre diet of the Carnivora (Chao *et al*., 2005; Vincze *et al*., 2022). Cheetahs have also been shown to have relatively high percentages of malignancy compared to other species with equivalent body mass, lifespan and litter size (Boddy *et al*., 2020). Although it is beyond the scope of this study to functionally validate them, the pseudogenisation events we identify are in a range of genes associated with important defects such as these, and therefore may contribute to conservation-relevant traits associated with health and fitness.

### PTCs shared between wild and captive cheetahs

Of the 22 genes with PTCs shared between multiple wild cheetahs, four were also identified in the chromosome-level cheetah assembly (VMU_Ajub_asm_v1.0, GCA_027475565.2): ARL13A, CFAP119, DEFB116 and NT5DC4.

The function of NT5DC4 (5’-Nucleotidase Domain Containing 4) is not well known, although it is part of the family that catalyses the intracellular hydrolysis of nucleotides. The related gene NT5C2 has been associated with immunological and metabolic disorders in humans <u>(Jordheim, 2018)</u>.

ARL13A (ADP Ribosylation Factor Like GTPase 13A) is predicted to be involved in ciliary structure and signalling <u>(Song and Perkins, 2018)</u>. In mice and humans, there is evidence that defects in ciliary motility and structure may cause respiratory infections and poor fertility (Tilley *et al*., 2015; Sironen *et al*., 2020). Other genes in the ARL family have been associated with fertility; ARL4 is linked to significantly reduced sperm count in mice (Schürmann *et al*., 2002).

Poor fertility is a widely-reported issue in both captive and wild cheetahs; male cheetahs have low sperm concentrations and high proportions of malformed sperm (Wildt *et al*., 1983; Lindburg *et al*., 1993; Crosier *et al*., 2007; Koester *et al*., 2015). In addition to ARL13A, CFAP119 (Cilia and Flagella Associated Protein 119, also known as Coiled-Coil Domain-Containing Protein 189 (CCDC189)) is also potentially involved in male fertility. Many CCDC genes have previously been linked to male fertility in mice and humans (Tang *et al*., 2017; Zhang *et al*., 2019), and a deleterious mutation in CCDC39 has previously been identified in the cheetah (Samaha *et al*., 2021).

Notably, a splice donor variant in CFAP119 was associated with asthenospermia (low sperm motility) in Nordic Red dairy cattle (Iso-Touru *et al*., 2019). The variant resulted in a frameshift mutation and premature translation termination, with the protein being truncated by over 40%, resulting in a lack of flagellar C1a complex subunit C1a-32, which modulates physiological movement of sperm flagella (Iso-Touru *et al*., 2019). A second coiled coil domain is found downstream of this, in exon 9 in the cattle. The novel PTC we identify in the cheetah may therefore cause a similar asthenospermic effect, as the cheetah protein is also truncated by 40% and the second coiled coil domain is likely lost. Further experimental validation is necessary to verify whether the PTC we observe in the cheetah impacts the protein in the same way.

Although the function of DEFB116 (Defensin Beta 116) is not known, other genes in the DEFB family are involved in antimicrobial defence in the skin and respiratory tract (Schröder and Harder, 1999; Schneider *et al*., 2005; Dhople *et al*., 2006) and regulation of sperm function (Zhang *et al*., 2018). Nearly all DEFB genes are preferentially expressed in the male reproductive tract and expression is enhanced during sexual maturation (Patil *et al*., 2005). In particular, DEFB29 is involved in sperm motility in mice and humans (Caballero-Campo *et al*., 2014) whilst DEFB23, DEFB26, and DEFB42 are linked with sperm motility and maturation in the epididymis of rats (Zhang *et al*., 2018).

The identification of the same PTC in more than one unrelated cheetah presents strong evidence that the mutations we identify are segregating across the African wild cheetah population. However, only 4 of the 22 PTCs supported by the population data are also observed in the long-read reference genome (VMU_Ajub_asm_v1.0, GCA_027475565.2). The population data we utilise (Dobrynin *et al*., 2015) and the original cheetah reference genome (aciJub1, GCA_001443585.1) are derived from individuals from Namibia and Tanzania, whilst VMU_Ajub_asm_v1.0 is derived from a captive individual (Lisbon Zoo) with at least 4 generations of controlled breeding in captivity (Marker and Johnston, 2022). The difference in occurrence of PTCs between the wild African cheetahs and the captive-bred individual highlights the importance of further population resequencing. If important and functionally-relevant pseudogenisation events are being overlooked due to a lack of sequence data, this has the potential to limit effective design for captive breeding programmes and could potentially lead to wider fixation of deleterious mutations in this endangered species.

## Conclusion

The cheetah has experienced severe genetic bottlenecks and a prolonged low effective population size for millions of years. The resulting accumulation of weakly deleterious mutations across the genome is likely to have contributed to the prevalence of diseases and disorders in both captive and wild cheetahs today. The growing effort to generate high-quality reference genomes can contribute to comparative genomic investigations of species-specific gene family dynamics and pseudogenization. Here, we identified 89 genes with novel premature stop codons resulting in gene pseudogenization. Of these, at least 22 are shared in wild cheetahs and four are observed in both captive and wild cheetahs. The four genes with potentially fixed PTCs across the species are involved in fertility and immune response and may be contributing to the reproductive and immune defects observed in cheetahs.

## Acknowledgements

This work was supported by the UKRI Biotechnology and Biological Sciences Research Council Norwich Research Park Biosciences Doctoral Training Partnership [BB/T008717/1] and BBS/E/ER/230001A, BBS/E/ER/230001B, BBS/E/ER/230001C, BBS/E/ER/230002A and BBS/E/ER/230002B. Authors also acknowledge support from BBSRC Core Capability Grant BB/CCG1720/1 and the work delivered via the Scientific Computing group, as well as support for the physical HPC infrastructure and data centre delivered via the NBI Research Computing group. The authors would like to thank Anil S. Thanki for his guidance running GeneSeqToFamily.

## Author contribution statement

Language used to describe roles below uses the CRediT Taxonomy (credit.niso.org).

Conceptualization: WH; data curation: WN, WH, formal analysis: JP, WN, WH; funding acquisition: WH; investigation: JP, WN, WH; methodology: JP, WN, WH; project administration: JP, WN, WH, resources: WH; software: WN, WH, JP; supervision: WH, WN; validation: JP; visualization: JP; writing – original draft: JP; writing – review & editing: JP, WN, WH.

## Conflict of interest

The authors have no competing interests.

## Notes

### Competing Interest Statement

The authors have declared no competing interest.

